# Cortical selectivity driven by connectivity: Innate connectivity patterns of the visual word form area

**DOI:** 10.1101/712455

**Authors:** Jin Li, David E. Osher, Heather A. Hansen, Zeynep M. Saygin

**Affiliations:** Department of Psychology & Center for Cognitive and Brain Sciences, The Ohio State University, Columbus, OH

## Abstract

What determines the functional organization of cortex? One hypothesis is that innate connectivity patterns set up a scaffold upon which functional specialization can later take place. We tested this hypothesis by asking whether the visual word form area (VWFA), an experience-driven region, was already connected to proto language networks in neonates scanned within one week of birth. With resting-state fMRI, we found that neonates showed adult-like functional connectivity, and observed that i) language regions connected more strongly with the putative VWFA than other adjacent ventral visual regions that also show foveal bias, and ii) the VWFA connected more strongly with frontotemporal language regions than with regions adjacent to these language regions. These data suggest that the location of the VWFA is earmarked at birth due to its connectivity with the language network, providing evidence that innate connectivity instructs the later refinement of cortex.

## INTRODUCTION

Decades of research suggest that the adult brain is composed of patches of cortex that are specialized for unique mental functions. To what extent is the functional organization of the human brain innate? Recent advances in developmental neuroimaging have made it possible to start to answer this question. For example, a previous study showed category-selective responses in high-level visual cortex for faces and scenes in infants^1^. Further, research in congenitally blind individuals suggests that cortical selectivity for high-level visual categories may not require visual experience^2^. In addition to the early emergence of visual processing, a previous study also found a neural precursor of language processing in infants^3^. Specifically, they found brain activity in left superior temporal and angular gyri to human speech in 3-month-old infants. These studies support the protomap hypothesis, which suggests that early genetic instructions give rise to the mature functional areas of cortex. However, the mechanisms that drive this early functional specialization remains ambiguous.

One possibility is that in the case of visual areas, pre-existing retinotopic biases may predispose a region to become selective to foveal or peripheral stimuli (Retinotopic/Eccentricity Hypothesis)^4–8^. However, this hypothesis is unlikely to fully explain the dissociation between face and word representations in visual cortex; faces and words are both foveal stimuli^8–10^ but are represented in different cortical areas. The Connectivity Hypothesis (which is not mutually exclusive from the Retinotopic Hypothesis) proposes that the specialization of a given brain region is largely shaped by how it connects and communicates with the rest of the brain. Previous work showed that structural connectivity (via diffusion imaging) as well as functional connectivity (via resting-state scans) can predict task-based selectivity across the brain^11, 12^. Further, Barttfeld et al. (2018) found a lateral-to-mesial organization in ventral visual cortex in newborns, suggesting that functional connectivity present at birth may constrain the subsequent functional specialization of visual areas^13^. This work suggests that connectivity is tightly intertwined with functional selectivity, and that perhaps early connectivity patterns may earmark the location of functionally selective cortices. Can the visual word form area (VWFA), which responds strongly to visual words or letter strings in literate individuals^14, 15^ be differentiated from the adjacent fusiform face area (FFA) by its connections to high-level cortex like the frontotemporal language network?

In adults, the VWFA connects with perisylvian language cortex, differentiating it from adjacent visual cortex^16^; other studies also found that white matter fibers that originated from the VWFA pass through fascicles that may be critical for language processing^17, 18^. In children, a longitudinal study found that connectivity patterns in pre-literate 5-year-olds predicted the location of the VWFA in each child at age 8 after they learned to read, and differentiated it from the adjacent FFA^19^. The connectivity patterns that predicted the VWFA included putative language areas, suggesting that connectivity to these regions may earmark the future location of the VWFA, and also set up a scaffold upon which future functional specialization can take place. However, while the 5-year-olds could not read (and at that age, lacked neural selectivity to letters or letter-like stimuli), they still would have had years of visual experience with letters and words. Is the putative VWFA already connected differently and set up to be differentiated from adjacent visual regions, even at birth with no visual experience with words and little visual experience at all?

Here, we tested this proto-organization of the VWFA in the newborn brain. Based on the Connectivity Hypothesis, we hypothesized that although the VWFA is highly experience-dependent, it is already ‘prewired’ to be selective for visual words by communicating with proto language regions at birth. By examining neonates who were scanned within one week of birth, we asked i) Do language regions show stronger functional connectivity (FC) with the putative VWFA than with other high-level visual areas like face, scene, and object areas? and ii) Does the VWFA show stronger FC with language regions than with adjacent frontotemporal regions like the multiple-demand (MD) network, speech regions, and primary auditory cortex (A1)?

## RESULTS

We examined whether the putative VWFA showed privileged connections with language regions even at birth. Because neonates cannot participate in task-based fMRI experiments, and because they do not yet have a VWFA, we overlaid functional parcels from previous studies and atlases^19–22^ to the neonates and adults in this study (see Online Methods for details). As an empirical check, we applied these parcels to an independent group of adults who performed the fMRI localizer tasks for all of the mental domains examined in the present study (see Supplementary Results 1&2 and Supplementary Figures 1&2). We successfully identified subject-specific functional regions (fROIs) within these parcels; these fROIs demonstrated functionally selective responses for the conditions of interest in independent fMRI runs (Supplementary Results 1&2 and Supplementary Figure 1&2). These results support the functional relevance and specificity of these parcels and the spatial variability of functional specialization across subjects which is captured by these parcels.

Note that all the analyses were performed on the volume and voxel-wise results were projected to the surface for a more comprehensive visual presentation of data.

### The selectivity of VWFA-language connections compared with other visual areas

First, we asked: do language regions selectively connect to the expected site of the VWFA, compared with other adjacent high-level visual regions? We compared the functional connectivity (FC) of language regions to the VWFA vs. to other high-level visual areas in the ventral stream, specifically in regions in the vicinity of the VWFA, including face selective regions (Fusiform Face Area, FFA; Occipital Face Area, OFA), scene selective region (Parahippocampal Place Area; PPA), and object selective regions (Lateral Occipital, LO; Posterior Fusiform Sulcus, PFS) (Fig. 1).

**Fig. 1.**
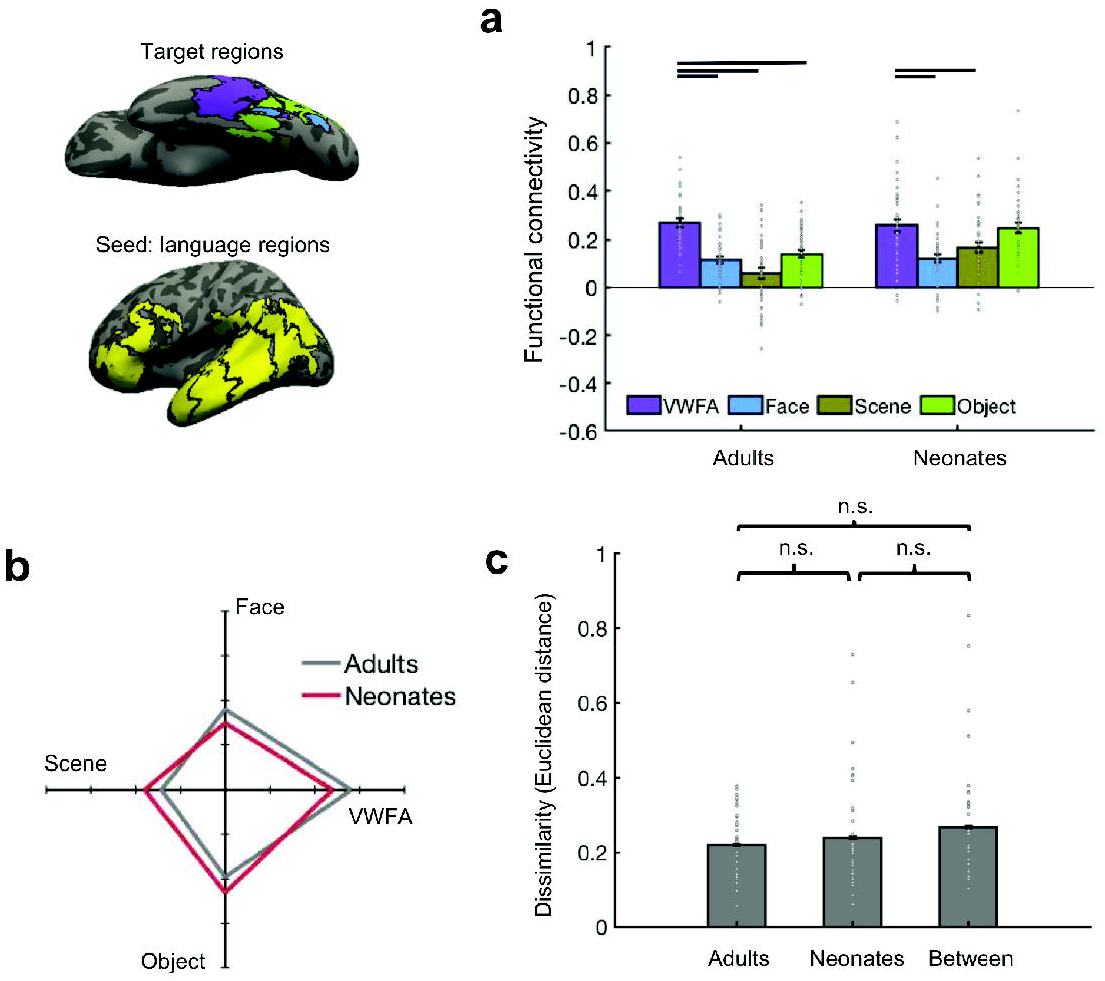
FC between language regions (seed) and high-level visual regions (targets). Seed, language (yellow); targets, VWFA (purple), faces (blue), scenes (olive), objects (light green). **(a)** Mean FC between language regions and high-level visual regions in ventral visual stream. Connectivity values were Fisher z transformed. Individual data points (n = 40 for each age group) were shown for each category. Error bars denote s.e.m. Horizontal bars reflect significant *post hoc* paired *t*-tests *p* < 0.05, corrected. **(b)** FC fingerprint of language regions. Connectivity values were mean-centered and averaged within each of the four categories to plot the relative patterns for the adult (n = 40) and neonate groups (n = 40). **(c)** FC pattern dissimilarity for within and between groups (n = 40 for each age group). Euclidean distance between each individual and others either from the same group or different group. n.s., non-significant.

We first performed a complete 2-way mixed design ANOVA with age group (neonate, adult) as the between-group variable and target (VWFA, faces, scenes, objects) as the within-group variable. We found significant main effects for both target and age group (target, (F(3,312) = 24.47, p < 0.001, partial η_2_ = 0.19, 95% CI of partial η_2_ = [0.11,0.26]; age group, F(1,312) = 14.07, p = 0.002, partial η_2_ = 0.04, 95% CI of partial η_2_ = [0.01,0.09]; and a significant interaction (F(3,312) = 4.92, p = 0.002, partial η_2_ = 0.05, 95% CI of partial η_2_ = [0.01,0.09]). Similar results were observed after accounting for size (Supplementary Results 3). *Post-hoc t*-tests revealed that in adults, language regions showed significantly higher FC with the VWFA than they did with faces (t(39) = 7.58, p<0.001), Cohen’s d = 1.20, corrected; 95% CI = [0.11,0.19]), scenes (t(39) = 9.39, p < 0.001, Cohen’s d = 1.49, corrected; 95% CI = [0.16, 0.25]) and objects (t(39) = 7.84, p < 0.001, Cohen’s d = 1.24, correct; 95% CI = [0.09, 0.16]) (Fig. 1b). The neonates showed a similar pattern, where connectivity between language regions and the VWFA was significantly higher than connectivity of language regions to face (t(39) = 6.28,p < 0.001, Cohen’s d = 0.99, corrected; 95% CI = [0.09, 0.18]) and scene (t(78) = 3.90, p < 0.001, Cohen’s d = 0.62, corrected, 95% CI = [0.04, 0.14]) regions, but we found no statistically significant evidence for a difference between language regions’ connectivity to the VWFA vs. object regions in neonates (t(39) = 0.55, p = 0.59, Cohen’s d = 0.09; 95% CI = [−0.03, 0.06]) (Fig. 1a). The same results were found when we intersected the functional parcels used here with meta-analysis maps generated from Neurosynth (https://neurosynth.org/; see Supplementary Results 5 and Supplementary Figure 6), suggesting that the results are largely applicable to other ways of defining functional brain regions and with narrower definitions of the functional parcels.

An exploratory analysis revealed that the VWFA was more connected than object regions to the more canonical aspects of the language network, the language parcel that likely encompasses Broca’s and the language parcel that like encompasses Wernicke’s areas (Online Methods) in neonates as well as in adults (neonates: Broca: t(39) = 3.06, p = 0.004, Cohen’s d = 0.48, corrected; 95% CI = [0.03, 0.15]; Wernicke: t(39) = 3.23, p = 0.003, Cohen’s d = 0.51, corrected; 95% CI = [0.04, 0.16]; adults: Broca: t(39) = 7.21, p < 0.001, Cohen’s d = 1.14, corrected; 95% CI = [0.21,0.38]; Wernicke: t(39) = 3.13, p = 0.003, Cohen’s d = 0.50, corrected; 95% CI = [0.03, 0.15]).

To further compare connectivity patterns between groups, we next looked at the connectivity fingerprints of language regions to visual cortex in neonates and adults (Fig.1b). Here we plot the relative connectivity of language regions to each of the four target categories (VWFA, face, scene, object regions) as compared to the mean of all four categories. We found that neonates had a very similar shape of the connectivity fingerprints as adults, suggesting similar FC patterns between groups. We statistically quantify the similarity of FC patterns between adults and neonates using Euclidean distance (as a measure of similarity) of the 4-dimensional FC pattern between participants. No statistically significant evidence for a difference between the within-group similarity and between-group similarity was found (within-adults vs. within neonates: t(78) = −0.72, p = 0.47, Cohen’s d = 0.16; 95% CI = [−0.07, 0.03]; within-adults vs. neonates-adults: t(78) = −1.68, p = 0.10, Cohen’s d = 0.38; 95% CI = [−0.11,0.01]; within-neonates vs. neonates-adults: t(78) = −0.85, p = 0.40, Cohen’s d = 0.19; 95% CI = [−0.10, 0.04]) (Fig.1c; see Online Methods for more details).

These results indicate that neonates show an overall similar FC pattern as adults, with the highest connectivity between language regions and the VWFA. Interestingly, neonates show similar connectivity between language-VWFA and language-object regions for the language network as a whole, but show dissociations in VWFA vs. object connectivity to the more canonical aspects of the language network, suggesting that further developmental refinement of connectivity does occur, especially to specific aspects of the language circuit.

### Functional connectivity at a voxelwise level in ventral temporal cortex

Next, we applied a voxelwise approach to analyze the functional connectivity profiles of language regions at a more fine-grain level. This analysis allowed us to examine spatial specificity of language-VWFA connectivity, which would serve as a complement to the parcel-wise analyses. We used language regions as the seed and we looked within ventral temporal cortex (VTC) for voxels that connected most with these language regions. Fig. 2a shows the connectivity of VTC voxels to language regions in representative neonates and adults. Consistent with the previous parcel-wise analysis, the voxels that have highest connectivity to language regions were mostly located in the lateral portion of VTC, which is within the putative VWFA location. To quantitatively identify which functional regions these voxels belonged to, we parametrically increased the connectivity threshold from the median to the top 95^th^ percentile of FC across VTC, and calculated the number of voxels within the VTC that were connected to language regions; we then quantified how many of these voxels belonged in each functional region using Dice coefficient (Online Methods). We found that voxels that were connected to language regions were always located in the expected VWFA, vs. all other functional regions in the vicinity; this result was significant for all thresholds (Fig. 2b).

**Fig. 2.**
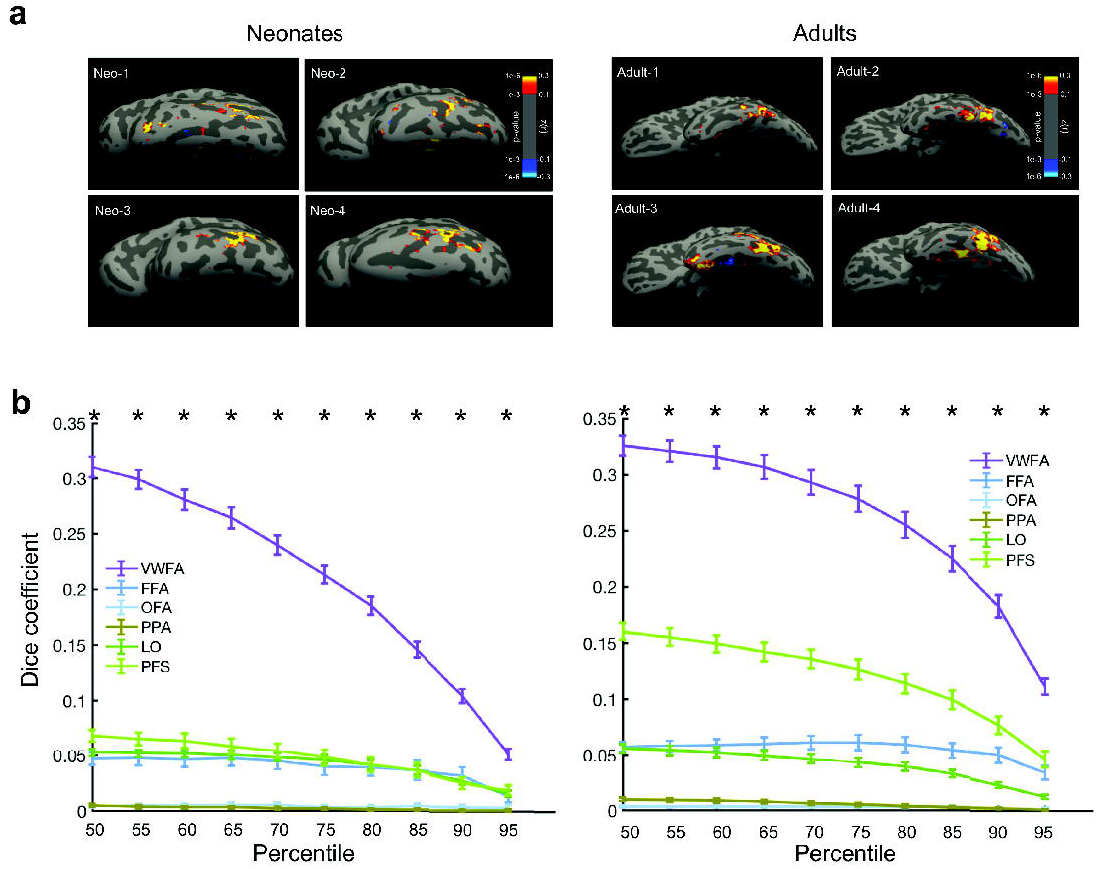
Voxel-wise analyses within the ventral temporal cortex (VTC) using language regions as the seed. **(a)** Heatmaps for voxels with connectivity to language regions in representative neonates and adults, thresholded at z(r) greater than 0.1 (p < 0.001). **(b)** Parametrically increasing the threshold of FC from the median to the 95^th^ percentile within VTC, we quantified how many of these voxels belonged in each functional region using *Dice coefficient*. Averaged FC (Fisher’s z transformed) across neonates (n = 40; 50th: z(r) = 0.25, p < 0.001; 95^th^: z(r) = 0.52, p < 0.001; Average FC across adults (n = 40; 50th: z(r) = 0.22, p < 0.001; 95^th^: z(r) = 0.45, p < 0.001). Error bars denote s.e.m across participants. * denotes significant paired *t*-test (VWFA vs. average of other functional regions, p < 0.05, corrected).

Overall, the parcel-based and voxelwise results indicate that the cortical tissue that may later develop sensitivity to visual words has connectivity patterns that are relatively adult-like in the neonatal brain, suggesting that it may be earmarked for function due to its preferential connectivity with language regions at birth. However, we also found differences between neonates and adults, especially with respect to object cortex, suggesting that there exist changes in this connectivity scaffold that likely result due to experience with literacy.

### Functional connectivity between the putative VWFA and language regions

Next, we asked, does the VWFA connect more to language regions vs. regions in the vicinity of language areas? We calculated FC between the VWFA (seed region) and the language, MD, speech, and A1 regions (target regions) (Fig. 3). We first performed a complete 2-way mixed design ANOVA with age group (neonate, adult) as the between-group variable and target (language, MD, speech, A1) as the within-group variable to examine VWFA’s connectivity. We found both that main effect of age and target were significant (age, F(1,312) = 9.29, p = 0.002, partial η_2_ = 0.03, 95% CI of partial η_2_ = [0, 0,07]; target, F(3,312) = 24.45, p < 0.001, partial η_2_ = 0.19, 95% CI of partial η_2_ = [0.11,0.26]), and the interaction was also significant (F(3,312) = 3.90, p = 0.009, partial η_2_ = 0.04, 95% CI of partial η_2_ = [0, 0.08]). Similar results were observed after accounting for size (Supplementary Results 3). Post-hoc t-tests revealed that in both adults and neonates the putative VWFA was more connected with language regions than with the other regions (Adults, MD: t(39) = 5.72, p < 0.001, Cohen’s d = 0.90, corrected; 95% CI = [0.10, 0.21]; Speech: t(39) = 6.48, p < 0.001, Cohen’s d = 1.02, corrected; 95% CI = [0.14, 0.26]; A1: t(39) = 9.32, p < 0.001, Cohen’s d = 1.47, corrected; 95% CI = [0.23, 0.36]; Neonates, MD: t(39) = 8.47, p < 0.001, Cohen’s d = 1.34, corrected; 95% CI = [0.11,0.18]; Speech: t(39) = 4.79, p = 0.028, Cohen’s d = 0.76, corrected; 95% CI = [0.06, 0.14]; A1: t(78) = 5.63, p < 0.001, Cohen’s d = 0.89, corrected; 95% CI = [0.09, 0.19]) (Fig. 3a).

**Fig. 3.**
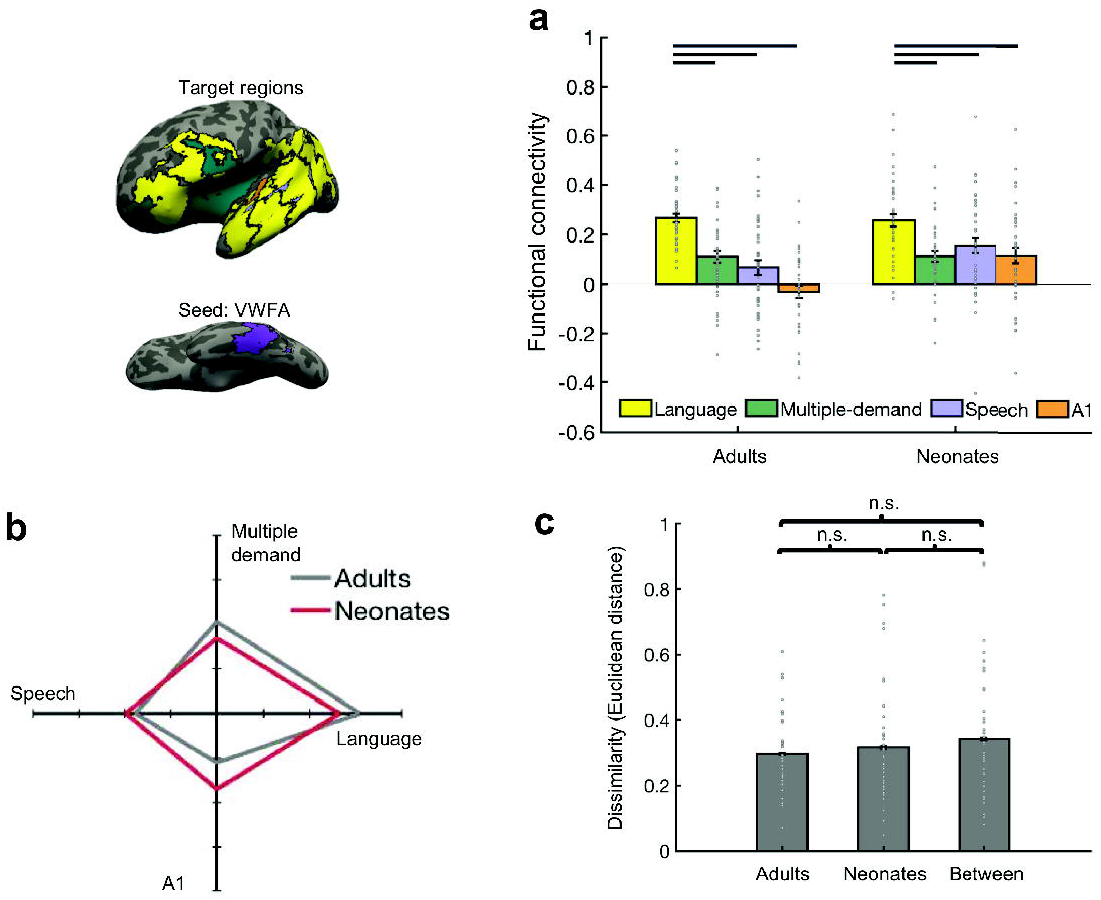
FC between VWFA (seed) and non-visual regions (targets). Seed, VWFA (purple); targets, language (yellow), speech (light purple), A1 (orange), MD (green). **(a)** Mean FC between VWFA and regions in temporal and frontal cortices. Connectivity values were Fisher z transformed. Error bars denote s.e.m. Individual data points (n = 40 for each age group) were shown for each category. Horizontal bars reflect significant *post hoc* paired *t*-tests *p* < 0.05, corrected. (**b**): FC fingerprint of VWFA. Connectivity values were mean-centered and averaged within each of the four categories to plot the relative patterns for the adult (n = 40) and neonate (n = 40) groups. **(c)** FC pattern dissimilarity for within and between groups (n = 40 for each age group). Euclidean distance between each individual and others either from the same group or different group. n.s., non-significant.

To further compare FC patterns between groups, we next plotted the connectivity fingerprint of the VWFA in neonates and adults, and observed similar fingerprint shapes between the two groups (Fig. 3b). Quantitative analyses of the similarity of FC profiles also confirmed this observation: no statistically significant evidence for a difference between the within-group similarity and between-group similarity was found (within-adults vs. within neonates: t(78)=-0.55, p=0.58, Cohen’s d = 0.12; 95% CI = [−0.08, 0.05]; within-adults vs. neonates-adults: t(78)=-1.29, p=0.20, Cohen’s d = 0.29; 95% CI = [−0.11,0.02]; within-neonates vs. neonates-adults: t(78)=-0.66, p=0.51, Cohen’s d = 0.15; 95% CI = [−0.11,0.05]) (Fig. 3c; see Online Methods for details).

To control for potential distance confounds and to further hone in on more spatially specific FC, we split up the target regions into frontal and temporal cortices and examined VWFA connectivity to language regions vs. adjacent functional regions in frontal and temporal cortex separately via planned t-tests. In both adults and neonates, the VWFA showed significantly higher FC to temporal language regions than adjacent speech (adults: t(39) = 5.48, p < 0.001, Cohen’s d = 0.87, corrected; 95% CI = [0.09, 0.20]; neonates: t(39) = 6.22, p < 0.001, Cohen’s d = 0.98, corrected; 95% CI = [0.08, 0.16]) and A1(adults: t(39) = 7.16, p <0.001, Cohen’s d = 1.13, corrected; 95% CI = [0.17, 0.31]; neonates: t(39) = 6.01, p < 0.001, Cohen’s d = 0.95, corrected; 95% CI = [0.11,0.22]) (Supplementary Figure 3a). Next, we compared the VWFA’s connectivity to frontal language regions vs. adjacent multiple-demand (MD) regions. In both adults and neonates, the VWFA showed significantly higher FC to frontal language regions as compared to adjacent MD regions (adults: t(39) = 9.71, p < 0.001, Cohen’s d = 1.54, corrected; 95% CI = [0.23, 0.35]; neonates: t(39) = 4.51, p < 0.001, Cohen’s = 0.71, corrected; 95% CI = [0.05, 0.13]) (Supplementary Figure 3b). The same results were found with Neurosynth-overlapped regions (Supplementary Figure 6).

Altogether, these results are consistent with previous adult studies by showing that the VWFA has higher FC to language related regions than to adjacent regions; here we also find that the neonatal VWFA has similar patterns of FC to language regions as the adult VWFA.

### Functional connectivity at a voxelwise level in frontal and temporal cortices

We also performed voxelwise parametric analyses for the frontal and temporal cortex using the VWFA as the seed. Consistent with the parcel-based analysis, we found that the voxels connected to the VWFA were most likely located in the expected language regions in temporal and frontal cortex in both neonates and adults (Supplementary Figure 4a and 4b). Heatmaps that illustrate the connectivity of frontotemporal voxels to the VWFA in representative neonates and adults are provided in Supplementary Figure 4c, and a volumetric presentation of average VWFA FC to the whole-brain in adults and neonates is also provided (Supplementary Figure 5). These results indicate that voxels connected to the VWFA are located within putative frontal and temporal language regions in neonates and adults alike.

## Discussion

A mosaic-like functional organization is consistently found in the adult brain. However, the driving factor of this functional organization and its variation across individuals remains unclear. The Connectivity Hypothesis proposes that the future function of a given brain area is largely shaped by how this region connects with the rest of the brain. Classic studies of ‘rewired’ ferrets showed that the cortical region that would have developed into A1 took on many of the properties of V1 after retinal input was rerouted to that location, showing in animal models that connectivity precedes function^23–27^. Here, we tested the Connectivity Hypothesis in human neonates and specifically for a high-level visual function that is uniquely human. We asked: is the putative VWFA already pre-wired at birth to develop differential functional specialization from its neighbors?

The VWFA serves as a good model to study the emergence of functionally selective regions since this region is highly experience-dependent. We first found higher connectivity of language regions with the VWFA than with adjacent regions in visual cortex, and we further replicated previous FC findings in adults^28^, showing higher connectivity of the VWFA with language regions that might be involved in different aspects of language processing (i.e., lexico-semantic processing, syntactic processing, structural processing) than with adjacent regions in frontotemporal cortex. Importantly, we also found that this region already shows adult-like connectivity patterns in neonates, suggesting that it may be earmarked to become selective to visual words by showing preferential connectivity with language regions. This research provides the earliest possible evidence in humans that the cortical tissue that will likely later develop sensitivity to visual words has a connectivity pattern at birth that makes it a fertile ground for such development – even before any exposure to words.

The organization of visual cortex, including high-level cortex, is largely biased by retinotopy^8–10^. This retinotopic organization is present very early in development, as evidenced by previous work, a recent study found that infant macaques, much like adults, showed a proto-organization for retinotopy throughout the visual system^29^. It is possible that early genetic instructions and underlying molecular/cytoarchitectonic determine the retinotopic preferences of neurons within these visual regions, including in high-level regions. Indeed, it has been posited that the VWFA starts out as part of the face network, and becomes increasingly selective to words and less selective to faces in the left hemisphere as literacy is acquired^15, 30^. This hypothesis is attractive because the perception of both faces and words require high-spatial frequency information that is represented foveally. Thus, with a retinotopic bias/connectivity from lower-level visual regions, it may be possible to first differentiate face regions from scene regions (foveal vs. peripheral bias) early in development (if not at birth), and then face from word regions after literacy is gained, perhaps through differential connections with fronto-temporal language regions. However, retinotopic organization or connectivity to early retinotopic cortex alone cannot explain the early differentiation of the VWFA from face regions, as we found here. We propose that in addition to its predisposition to foveal stimuli, the location of the future VWFA also depends on its innate connectivity with language regions even at birth.

Numerous open questions remain. First, what are the changes that occur with reading? In other words, if the VWFA is not selective to words in neonates, how does it become word-selective with literacy? Our results also suggest that there are certain differences between neonates and adults, such as overall differences in connectivity magnitude, strengthening of specific connections (i.e., increased connectivity between VWFA and frontal language regions in adults), and refinement of connections (i.e., decreased connectivity between VWFA and speech, A1 in adults). For example, our parcel-based analyses show that object regions also have high FC to language regions in neonates whereas in adults they do not; on a finer grain, our parametric analyses show that the putative VWFA is in fact differentiated from object regions in their connectivity to language cortex. Further, we found that the VWFA is also differentiated from object cortex by their connections to a narrower definition of language regions (i.e. putative Broca’s and Wernicke’s areas). These findings suggest that while the VWFA is already differentiated from face, scene, and to some extent object regions at birth, there is likely further refinement to fully differentiate orthographic representations from other objects in visual cortex, in line with e.g. Augustin et al. (2015)^31^ and Kubota et al. (2018)^32^. Experience with spoken and written language will likely strengthen connections with specific aspects of the language circuit and further differentiate this region’s function from its neighbors as an individual gains literacy.

Another question that remains unanswered is how the connectivity patterns themselves arose prenatally and evolutionarily. It is likely that a complex mechanism of intrinsic properties of cortical regions and early signaling mechanisms set up these large-scale connections. The VWFA may simply be in a privileged location, due to a myriad of mechanisms including cellular properties and intrinsic circuitry, in addition to large-scale connectivity that facilitates its later selectivity. Future studies combining animal models with studies in other human populations, e.g. premature human infants, may help further elucidate the evolution of these mechanisms. Moreover, the present study focused on functional connectivity, which raises another interesting question about whether there exists innate structural connectivity between the putative VWFA and language regions at birth and what its developmental trajectory looks like. A recent study observed white matter (i.e., arcuate fasciculus) alterations in 18 months infants with familiar risk of developmental dyslexia^33^. A potential future direction in this line of research is to explore the role of white matter maturation properties (e.g., fiber density and myelination) in prolonged language development, and examine how their interplay with functional connectivity and experience. Other open avenues of future research include looking at effective connectivity to try to tease apart the directionality of connectivity (which would need to be verified with animal models) as well as graph theoretical approaches^34^ to show similarities or differences in network structure between neonates and adults.

Finally, it remains unknown how laterality arises in the human brain. The questions and hypotheses of this study pertain to the canonical location of the VWFA within the context of the ventral visual stream (i.e. with respect to other category-selective regions), rather than the laterality bias of the VWFA vs. e.g. the FFA. Consistent with previous studies in adults that compared VWFA connectivity to adjacent regions^16, 28, 35^, we restricted our analyses to the left hemisphere in order to eliminate potential confounds of cross-hemispheric differences. The development of functional specificity in right vs. left visual cortex remains an open question. A longitudinal study that can functionally define these regions in children (after they can participate in a task-based fMRI scan) can answer these questions and more, as discussed in the limitations below.

There exist certain limitations in the current study. We found evidence in favor of the Connectivity Hypothesis; stronger causal evidence would involve experimental manipulations of connectivity patterns to test if functional specialization changes as a consequence of these connectivity changes. However, this type of study would be invasive in newborn humans; here we attempt to leverage experience-dependent domains and a study of neonates to test the Connectivity Hypothesis but acknowledge the limitations of causal inferences that can be drawn from noninvasive studies. Further, a challenge of studying the functional organization of the neonatal brain is that there is no adequate way to localize functional responses using fMRI in neonates. Here we used functional parcels from previous studies and overlaid these parcels onto both adult and neonate brains. These parcels likely encompass the functional regions in individual subjects which offer better functional relevance than anatomical landmarks, but as a consequence, also likely overestimate the size of the functional regions. To further explore spatial specificity, we chose adjacent functional parcels as comparisons, explored smaller subsets of the language parcels, and performed voxelwise analyses on individual subject data without predefined functional regions. However, the present results are still limited by the functional parcels as well as current registration and image processing methods in neonates; better registration methods such as surface-based registration to an adult template are currently unavailable in neonates but will likely improve the results and inferences drawn from these studies. Additionally, future studies may consider new approaches to localize functional responses in young infants or longitudinal studies, e.g., Saygin et al. (2016)^18^, to define each of these functional regions in individual subjects and further test the specificity of the current findings. Finally, we tested the Connectivity Hypothesis for the VWFA specifically. The findings suggest that connectivity-based scaffolding may be a general driving mechanism for the functional organization of human cortex, but the generality of this hypothesis for other mental domains remains to be tested.

## Online Methods

### Participants

#### Neonates

We used the initial release of the Developing Human Connectome Project (dHCP) neonatal data (http://www.developingconnectome.org)^36^. Neonates were recruited and imaged at the Evelina Neonatal Imaging Centre, London. Informed parental consent was obtained for imaging and data release, and the study was approved by the UK Health Research Authority. All 40 neonates of the initial release were included in functional connectivity analysis and were born and imaged at term age (15 female, mean gestational age at birth = 38.99 weeks, gestational age range at scan = 37-44 weeks).

#### Adults

Adult data were obtained from the Human Connectome Project (HCP), WU-Minn HCP 1200 Subjects Data Release (https://www.humanconnectome.org/study/hcp-young-adult)^37^. All participants were scanned at Washington University in St. Louis (WashU). 40 adults were included in functional connectivity analysis (15 female, age range = 22-36 years old). These adult participants were motion and sex matched to the neonates. Specifically, for each neonatal participant we matched with an adult from the HCP dataset with the same sex who showed the most similar motion parameter (i.e., framewise displacement, FD) with the k-nearest neighbors’ approach. By doing this, we are able to match the sex ratio and no evidence for a statistically difference was found for head motion between groups (t(78) = 0.77, p = 0.45, Cohen’s d = 0.17, 95% CI = [−0.02, 0.01]).

### Data acquisition

#### Neonates

Imaging was carried out on 3T Philips Achieva (running modified R3.2.2 software) using a dedicated neonatal imaging system which included a neonatal 32 channel phased array head coil^38^. All neonates were scanned in natural sleep; previous studies have shown that the resting-state FC remains consistent while awake and asleep, as well as while under anesthesia^39, 40^.

##### Resting-state fMRI

High temporal resolution fMRI developed for neonates using multiband (MB) 9x accelerated echo-planar imaging was collected (TE/TR = 38/392ms, voxel size = 2.15 × 2.15 × 2.15mm^3^). The duration of resting-state fMRI scanning was approximately 15 minutes and consisted of 2300 volumes for each run. No in-plane acceleration or partial Fourier was used. Single-band reference scans were also acquired with bandwidth matched readout, along with additional spin-echo acquisitions with both AP/PA fold-over encoding directions.

##### Anatomical MRI

High-resolution T2-weighted and inversion recovery T1-weighted multi-slice fast spin-echo images were acquired with in-plane resolution 0.8 × 0.8mm^2^ and 1.6mm slices overlapped by 0.8mm (T2-weighted: TE/TR = 156/12000ms; T1 weighted: TE/TR/TI = 8.7/4795/1740ms).

#### Adults

All the scans of WU-Minn HCP 1200 Subjects Data Release was carried out using a customized 3T Connectome Scanner adapted from a Siemens Skyra (Siemens AG, Erlanger, Germany) with 32-channel Siemens receive head coil and a “body” transmission coil designed by Siemens specifically for the smaller space available using the special gradients for the WU-Minn and MGH-UCLA Connectome scanners.

##### Resting-state fMRI

Participants were scanned using the Gradient-echo EPI sequence (TE/TR = 33.1/720ms, flip angle = 52°, number of slices = 72, voxel size = 2 × 2 × 2 mm^3^). The duration of resting-state fMRI scanning was approximately 15 minutes and consisted of 1200 volumes for each run. All participants accomplished two resting-state fMRI sessions. Within each session, there were two phases encoding in a right-to-left (RL) direction in one run and phase encoding in a left-to-right (LR) direction in the other run. In current analysis, we used the LR phase encoding from the first session. Participants were instructed to open their eyes with relaxed fixation on a projected bright cross-hair on a dark background.

##### Anatomical MRI

High-resolution T2-weighted and T1-weighted images were acquired with isotropic voxel resolution of 0.7mm3 (T2-weighted 3D T2-SPACE scan: TE/TR = 565/3200ms; T1-weighted 3D MPRAGE: TE/TR/TI = 2.14/2400/1000ms)

### Preprocessing

#### Structural data Preprocessing

The dHCP data were released as preprocessed data; they used the dHCP structural minimal preprocessing pipeline^36^, briefly: bias correction, brain extraction using BET from FSL, and segmentation of the T2w volume using DRAW-EM algorithm^41^ which were developed for neonatal brain segmentation. Gray and white matter masks were obtained from segmentations using DRAW-EM algorithm provided by dHCP. The HCP data were released as preprocessed data; they used the HCP structural preprocessing pipeline^42^, briefly: gradient distortion correction, brain extraction, a bias field correction, and registration between the T2-weighted scan and T1-weighted scan. Each individual brain was also aligned to common MNI152 template (with 0.7mm isotropic resolution). Then, the FreeSurfer pipeline (based on FreeSurfer 5.3.0-HCP) was performed to segment the volume into predefined structures and surface reconstruction.

#### Functional data Preprocessing

The pre-processed functional data released by the dHCP had already undergone basic pre-processing steps (for details see Fitzgibbon et al. (2019)^43^): distortion-correction, motion correction, 2-stage registration of the MB-EPI functional image to T2 structural image and also generated a combined transform from MB-EPI to 40-week T2 template, and ICA denoising using ICA-FIX^44^. The data released by the HCP had already undergone basic pre-processing steps (for details see Glasser et al. (2013)^42^): removed spatial distortions, corrected for motion, registered the fMRI data to both structural and MNI152 template, reduced the bias field, and ICA denoising using ICA-FIX^44^. The HCP data were registered to each individual’s native space using the transformation supplied by the HCP and the following steps were performed on both the HCP and dHCP data: applied smoothing (Gaussian filter with the FWHM = 3 mm) within the all gray matter, and band-pass filter at 0.009-0.08 Hz. As a further denoising step, we used aCompCor44 to regress out signals from white matter and cerebrospinal fluid (CSF) to control physiological noise like respiration and heartbeat as well as non-neuronal contributions to the resting state signal. All the FC analyses were performed in native functional space.

### Defining the functional parcels

The parcels used here were originally created from probabilistic maps of functional activation across independent groups of participants, and are generated such that they encapsulate most individuals’ functional regions, via the group-constrained subject-specific method (GSS)^20^. Contrary to traditional group-based methods (e.g., random-effects analyses) or using anatomical approximations or Talairach coordinates based on meta analyses, the GSS approach takes individual variability of functional responses (size, shape, and location) into account, providing the anchor space for functionally specialized regions that activate systematically across individuals. The present study especially benefits from this approach due to the study of nonverbal neonates. Additionally, these GSS studies were chosen particularly because the tasks and fMRI contrasts that were used to define the functional regions of interest offer better controls for the domains of interest. All parcels are available online or via contacting the corresponding author of the cited publications.

All parcels were mapped to the FreeSurfer CVS average-35 MNI152 brain (if they were not already publicly provided in that space) and were subsequently registered to each individual’s brain (see below). <u>Language regions</u> were released by Fedorenko et al. and were defined by Sentences vs. pronounceable non-word sentences^20^ thus controlling for prosody, low-level auditory features, and speaker identify, and are found to respond similarly to auditory and visual versions of the stimuli^45, 46^. Temporal regions included: AntTemp, anterior temporal lobe; MidAntTemp, middle-anterior temporal lobe; MidPostTemp, middle-posterior temporal lobe; PostTemp, posterior temporal lobe; and AngG, angular gyrus. Frontal regions included: IFG, interior frontal gyrus; and IFGorb, orbital IFG. To get a narrower definition of language regions, we selected the IFG language parcel for Broca’s area and the MidAntTemp language parcel for Wernike’s area. The <u>speech region</u> was from Basilakos et al. (2018)^47^ and the region we used was in superior temporal gyrus, which was shown to be sensitive to the phonemic structure of human speech rather than low-level auditory properties or task-difficulty. <u>A1</u> was anatomically defined as Heschl’s gyrus (superior and transverse temporal cortex from the FreeSurfer Desikan-Killiany parcellation^48^ in CVS average-35 MNI152 space). <u>Multiple-demand (MD)</u> parcels located in left frontal cortex were obtained from Fedorenko et al. (2013)^21^, showing activation to hard vs. easy conditions of working memory tasks^21, 46, 49^. These parcels were in MFGorb, orbital part of the middle frontal gyrus; Insula; IFGop, opercular part of the inferior frontal gyrus; SMA, supplementary motor area; and ACC, anterior/mid cingulate cortex. The <u>VWFA</u>, located in left occipitotemporal cortex, was created from Words vs. line drawings of Objects, from Saygin et al. (2016)^18^. The other high-level visual parcels were derived from Julian et al.^22^, and were based on responses to dynamic movie clips^50^ and activation for the contrast of interest. <u>FFA and OFA</u> located in the fusiform and occipital cortex respectively were identified with faces > objects contrasts^51–53^; scene selective <u>PPA</u> was identified with scenes > objects contrast^54^ and was located in the parahippocampus; object selective <u>LO and PFS</u> were defined with objects > scrambled objects contrasts^55^ and located in the lateral occipital and posterior fusiform sulcus respectively. Because both VWFA and language are largely left lateralized^30, 56^, our study includes left hemisphere seeds and targets only, as was the case with previous studies of these regions^16, 28^ as well as a recent study which also looked at VWFA connectivity using the adult HCP dataset^35^.

All functional parcels were placed in the template CVS average-35 MNI152 space, and were overlaid onto each individual’s native anatomical brain using Advanced Normalization Tools (ANTs version 2.1.0; http://stnava.github.io/ANTs)^57–59^ for both adults and neonates. For registration between modalities (i.e., anatomical to native functional image for neonates), we used nearest neighbor interpolation with Freesurfer’s mri_vol2vol function (https://surfer.nmr.mgh.harvard.edu/fswiki/mri_vol2vol). To ensure no voxel belonged to more than one functional parcel, we assigned any intersecting voxels of two functional parcels to the one with smaller size as a way to compensate size differences (e.g., Brissenden et al. 2016^60^). Additionally, voxels within white matter and cerebellum were also removed. In total, we used 20 non-overlapping functional parcels from eight categories in the present study.

### Calculating functional connectivity

The mean timecourse of each functional parcel was computed from the preprocessed resting state images, and FC was calculated with Pearson’s correlation between the mean timecourse of each seed parcel and each target parcel. To generate normally distributed values, each FC value was Fisher z-transformed.

### FC fingerprint plots

First, we calculated the average FC from the seed to each of the target categories. Then we subtracted the mean FC across all categories from each of the averaged FC. Thus, the value in the fingerprint plots indicates how the seed connects to the targets compared to the mean connectivity of the seed to all categories (mean-centering) across subjects in each group. We further quantified the similarity of FC patterns between adults and neonates. Specifically, for each participant, the Euclidean distance was calculated between the 4-dimensional FC pattern of the seed (i.e., VWFA or language regions) and the average FC pattern of others either from the same group or the different group. This measured how similar each participant was to others.

### Voxel-wise FC analysis in the ventral temporal cortex (VTC) and frontotemporal cortex

We performed a voxel-wise analysis across VTC to get a finer characterization of the connectivity pattern with language regions. We defined the VTC from the Desikan-Killiany parcellation^48^, including the fusiform and inferior temporal labels, in FreeSurfer CVS average-35 MNI152 space, which were registered to each individual’s anatomy and masked with the gray matter image for each individual subject (as provided by the HCP and dHCP datasets). FC was computed between the mean timecourse of the language regions and the timecourse of each VTC voxel. Without predefining any functional parcels within the VTC, this analysis allowed us to characterize where the voxels with highest connectivity were located within the VTC. To quantify this, we performed a parametric analysis where we increased the threshold of FCs across all VTC voxels from the 50^th^ percentile (median) to the 95^th^ and calculated the overlap of these voxels with each of VTC regions with Dice coefficient. Specifically, each percentile determines the threshold for binarizing the connectivity data and overlap is calculated using Dice coefficient for each subject. For example, for the 50th percentile threshold, Dice coefficient was calculated by 2 * (A AND B) / A OR B, where set A are the voxels in VTC that are connected to language regions above the 50th percentile and set B are the voxels within the VWFA. We used Matlab to calculate percentiles and Dice coefficient. The same analysis was performed for frontal cortex and temporal cortex separately, and frontal and temporal cortex were again defined with Desikan-Killiany parcellation in the CVS average-35 MNI152 space and masked to only include gray matter within each subject’s individual space). Temporal cortex analyses were restricted to the more superior regions to prevent overlap with the VTC analysis. Individual subject results were projected to the surface of each subject using the surfaces provided by the dHCP and HCP with trilinear interpolation, which takes the average across the surface normal.

### Statistics

2-way mixed design ANOVA were used to test our main focus. Age group (adults, neonates) was the between-subject variable and target (i.e., different target categories) was our within-subject variable (i.e., repeated-measures), and thus there was no experimental group randomization or blinding in the present study. Paired *t-*tests were conducted for within group comparisons and two-tailed *t*-tests for across-group comparisons. The 95% confidence interval of the mean FC true population difference was also reported for each *post hoc t-*test. Benjamini & Hochberg/Yekutieli false discovery rate control (FDR)^61^ was used for multiple comparisons correction. Each post hoc t-test was corrected for the total number of paired-wise comparisons for each analysis. Data distribution was assumed to be normal, but this was not formally tested.

## Data availability

The data used in this study are publicly available. All relevant accession codes are publicly available in HCP (https://www.humanconnectome.org/study/hcp-young-adult) and dHCP (http://www.developingconnectome.org).

## Code availability

The code that supports the findings of this study are available from the corresponding author upon request.

## Acknowledgements

We would like to thank the Human Connectome Project (https://www.humanconnectome.org) and developing Human Connectome Project (http://www.developingconnectome.org). Analyses were completed using the Ohio Supercomputer (https://www.osc.edu). We would like to thank members of the Saygin Developmental Cognitive Neuroscience Lab for feedback and comments and for Drs. Fedorenko and Kanwisher for help with the localizers for the Supplementary Results. Funding: This research was partly funded by the Alfred P. Sloan Foundation (to Z.M.S) and Supplementary Results data were partly funded by NICHD/NIH grant F32HD079169 to Z.M.S. and MINT foundation grant to Kanwisher.

## Author contributions

J.L.: conceptualization, formal analysis, methodology, writing-original draft & editing; D.E.O. and H.A.H.: methodology, writing-review; Z.M.S conceptualization, supervision, writing-review & editing.

