## Supplementary Materials for "Cortical selectivity driven by connectivity: Innate connectivity patterns of the visual word form area"

**Supplementary Information**

Supplementary inventory:

**Supplementary Results 1&2:** Using four fMRI tasks, subject-specific functional regions (fROIs) were identified in an independent group of adult subjects within the functional parcels used in the main analyses. fROIs across subjects demonstrate variability in spatial location across subjects and demonstrate the need for larger parcel regions that will certainly encompass the sites of functional specificity in the neonates in the main study. Percent signal change was also extracted in independent runs to demonstrate functional specificity of these fROIs.

**Supplementary Results 3:** Two-way mixed design ANOVA of language regions’ FC using size as a covariate and two-way mixed design ANOVA of VWFA’s FC using size as a covariate.

**Supplementary Results 4**: FC between VWFA and temporal regions, and VWFA and frontal regions.

**Supplementary Results 5**: FC results for new Neurosynth-overlapped parcels.

**Supplementary Results 6**: Comparison between registration accuracy of ANTs and FLIRT.

**Supplementary Figure 1:** Probabilistic maps created based on fROIs across a group of adults (independent of those in the main study).

**Supplementary Figure 2:** Functional response profiles from separate runs than those used to define the fROIs in these independent adult subjects.

**Supplementary Figure 3:** FC between VWFA (seed) and temporal and frontal regions.

**Supplementary Figure 4:** Voxel-wise analyses from VWFA to frontotemporal cortices.

**Supplementary Figure 5:** Averaged whole-brain functional connectivity maps of VWFA for adults and neonates.

**Supplementary Figure 6:** FC between language (seed) and visual regions and FC between VWFA (seed) and temporal and frontal regions with new Neurosynth-overlapped parcels.

**Supplementary Figure 7:** Registration results.

**Supplementary Table 1:** Information of parcels used in the current study.

**Supplementary Results 1.** As part of a previous study conducted at MIT, an independent group of adults (N = 14, mean age=27.0 ±0.87, 6F; all right-handed) were recruited from the greater Boston area. Participants were screened for history of mental illness, gave written informed consent, and were compensated at $30 per hour. The study was approved by the Massachusetts Institute of Technology ethics committee. Participants were run on the VWFA, Language & Speech, MD, and Dynamic localizers, which are the same or similar fMRI localizers as those used in the previous studies from which we drew the functional parcels in the main analyses; see **Online Methods** and **Supplementary Table 1**. Experimental parameters were similar or identical to those used in the published studies for each localizer. Briefly, participants viewed the VWFA localizer where they saw words, line-drawings of faces, and objects (details in Saygin et al. 2016) and 3 additional fMRI localizers: the Language & Speech localizer with auditory stimuli consisting of English sentences, sentences with similar prosody but constructed from nonsense words, and statistically-matched textured speech which controlled for low-level auditory features but did not have recognizable speech sounds (Fedorenko et al., 2010; see also Overath et al., 2015; McDermott and Simoncelli, 2011); the MD localizer with Hard and Easy task conditions of a spatial working memory paradigm (Fedorenko et al. 2013); and the Dynamic localizer consisting of movie clips of faces, bodies, objects, and scenes (Pitcher et al., 2011; Julian et al. 2011). Data were collected on a 3T Siemens scanner with a 32-channel head-coil. Acquisition parameters were similar or identical to those used in the published studies for each localizer: the VWFA localizer was acquired with 2mm^3^ resolution, 2s TR, 30ms TE, 90° flip, 100x100 base resolution, 25 slices approximately parallel to the base of the temporal lobe to cover the entire inferior temporal cortex. We additionally collected a field map for distortion correction with the same slice prescription as the fMRI sequence (25 slices, 2mm^3^ resolution, 500ms TR, 55° flip, 100x100 base resolution). The Language & Speech and MD localizers were acquired with 2s TR, 30ms TE, 90° flip, GRAPPA acceleration factor 2, 2x2mm in-slice resolution, 4mm slice thickness, 31 slices for full-brain coverage; and the Dynamic localizer was acquired with 2TR, 30ms TE, 90° flip, 3mm^3^ voxels, and 32 slices for full-brain coverage. Experiments were counter-balanced across participants. A high-resolution (1 mm^3^) three-dimensional magnetization-prepared rapid acquisition with gradient echo (MPRAGE) scan was also acquired in all participants. All the imaging data were analyzed using standard pre-processing steps with Freesurfer ([www.surfer.nmr.mgh.harvard.edu/](http://www.surfer.nmr.mgh.harvard.edu/)) and FsFast ([www.surfer.nmr.mgh.harvard.edu/FsFast/](http://www.surfer.nmr.mgh.harvard.edu/FsFast/)). Images were motion-corrected, de-trended, and fit using a standard gamma function (d = 2.25 and t = 1.25). Runs were registered to each subject’s anatomical image using Freesurfer’s bbregister.

We applied the functional parcels used in the main analyses to identify subject-specific functional regions (fROIs) using the same GSS approach and contrasts as outlined in the **Online Methods**. Individual fROIs were defined using the top 10% most active voxels within these parcels for each contrast of interest: words > line-drawings of objects for the VWFA, line-drawings of faces > objects for face regions, English > nonword sentences for language regions, nonword sentences > texturized speech for speech region, hard > easy for MD regions, movie clips of scenes > objects for scene regions and objects > scrambled objects for object regions. Each subject’s fROI was registered to Freesurfer’s CVS_MNI152 standard space and probabilistic heat maps were created based on these adults (shown in **Supplementary Figure 1**). The figure shows that these functionally selective fROIs are somewhat consistent across participants, where more than half of subjects’ (>=7) fROIs land within similar spatial locations; however, we also observed large inter-subject variability as evidenced by the large spread of fROIs across e.g. temporal cortex for temporal language fROIs across subjects. These fROIs demonstrate variability in spatial location across subjects and demonstrate the need for larger parcel regions that will certainly encompass the sites of functional specificity in the neonates in the main study.

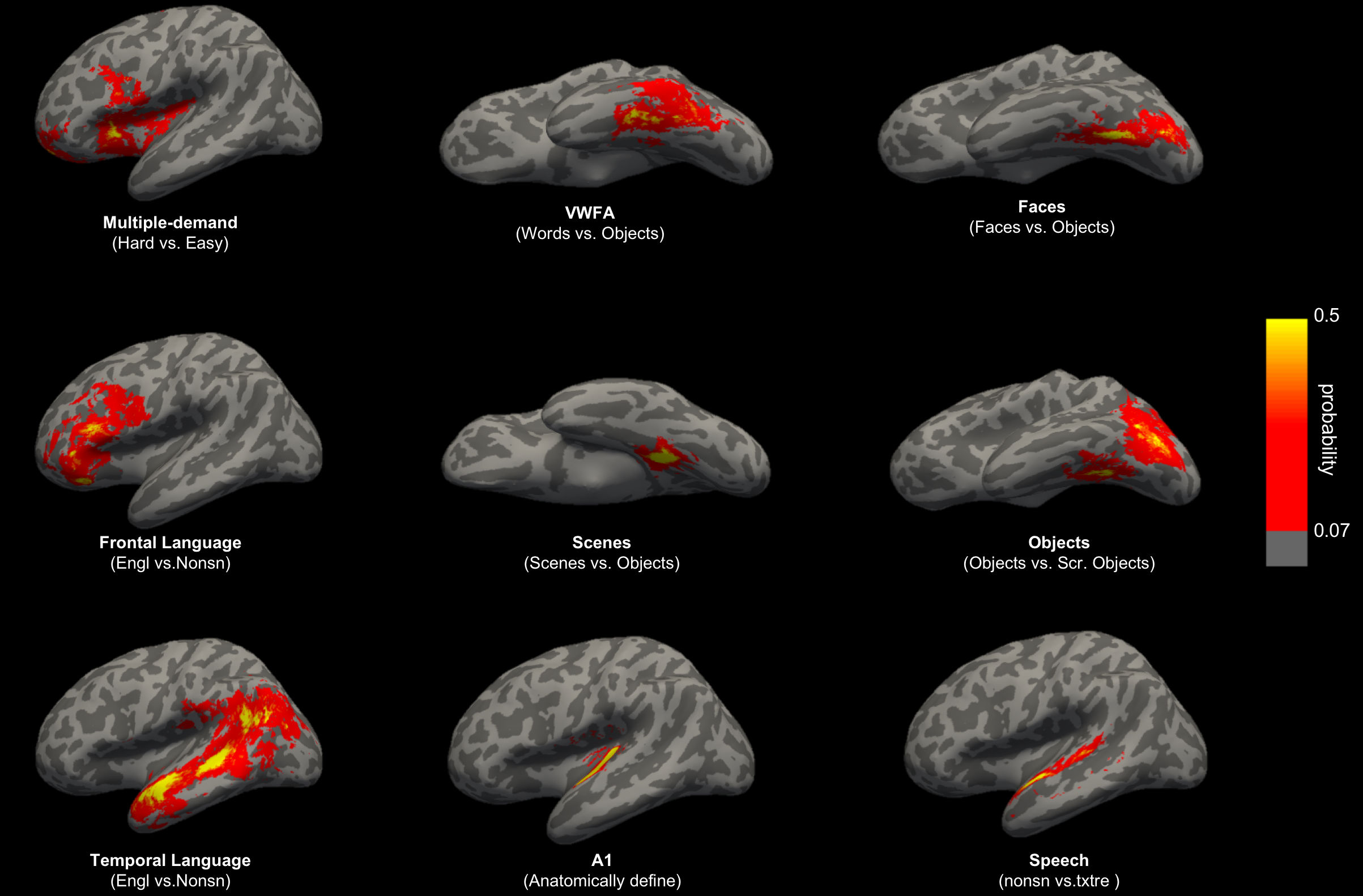

**Supplementary Figure 1 | Probabilistic maps created based on subject-specific functional regions.** Heatmaps show voxels that show functional selective response to a given contrast in at least 1 subject (prob. = 0.07) to equal or greater than 6 subjects (prob. = 0.5). Contrasts of interest for each region are provided in the text and below each region; note that A1 for each individual was anatomically defined in Heschl’s gyrus (superior and transverse temporal cortex from the FreeSurfer Desikan-Killiany parcellation). Engl, English sentences; Nonsn, non-word sentences; txtre, texturized; Src. Objects, scrambled objects. These fROIs demonstrate variability in spatial location across subjects and demonstrate the need for larger parcel regions that will certainly encompass the sites of functional specificity in the neonates in the main study.

**Supplementary Results 2.** Additionally, we extracted percent signal change (PSC, from independent fMRI runs from those used to define the fROIs) for the different experimental conditions across the different localizers to demonstrate functional selectivity of responses. fROI PSCs were averaged for each category of interest and nonparametric Wilcoxon sign-rank tests were performed to see if the conditions of interest for each fROI were higher than all of the other conditions (i.e. one test was performed for each fROI). **Supplementary Figure 2** shows functional response profiles for fROIs that were identified within the functional parcels, and demonstrates distinct response patterns across fROIs. With the exception of frontal lang and MD which showed trending effects, all other fROIs showed significantly higher responses to the mental function they are posited to be selective to vs. other conditions (p<0.05, Bonferroni-Holm corrected). Altogether, Supplementary Figures 1 and 2 suggest that the functional parcels used in the main results will capture both functionally specific responses to the functions of interest tested in the present study and also tolerate variability between individuals.

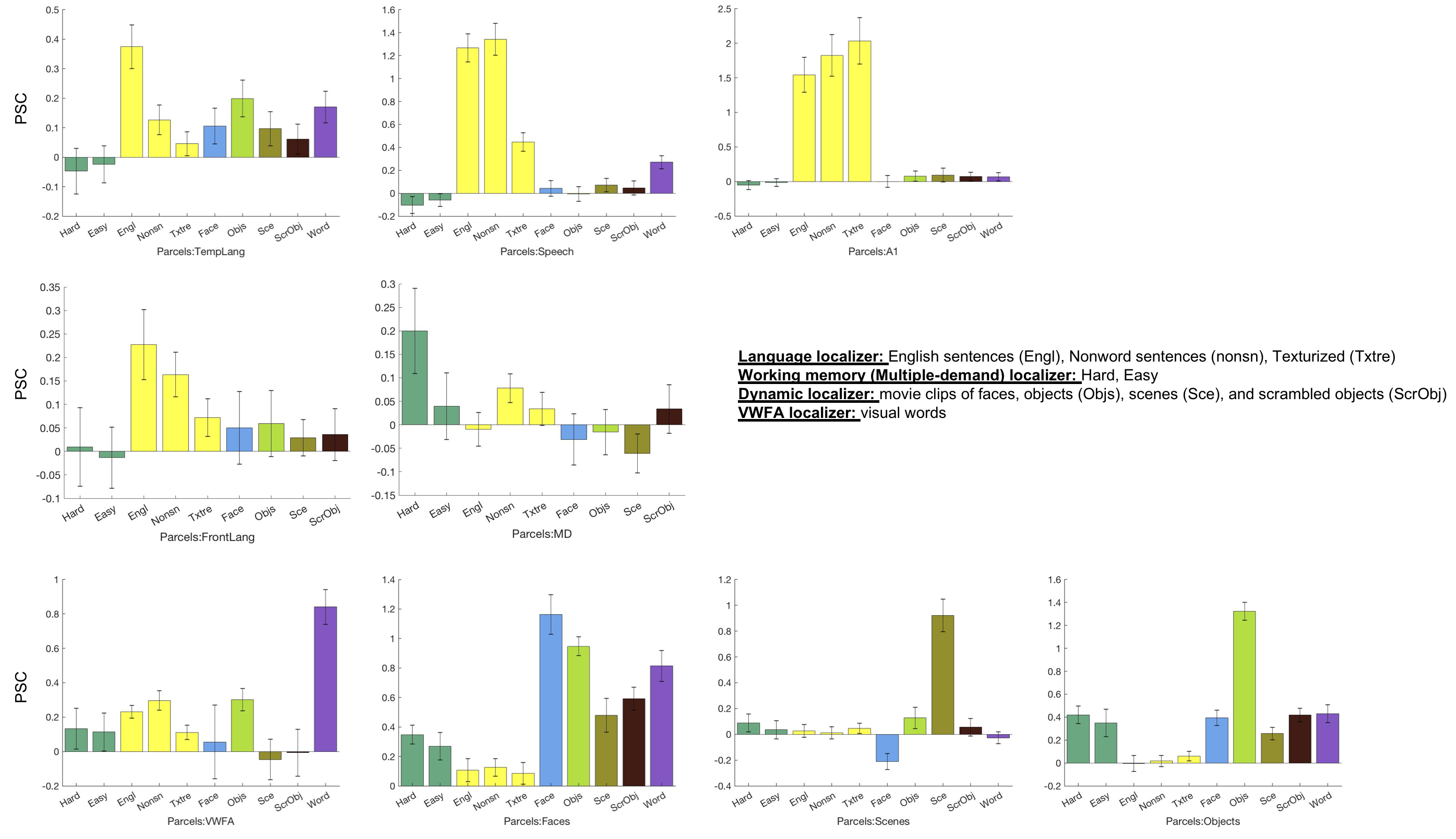

**Supplementary Figure 2 | Functional response profiles for every fROI.** Percent signal change (PSC) was extracted from independent run (separate from those used to define each region) for every functional localizer. Note that the field-of-view (FOV) of the VWFA localizer only had temporal cortex coverage so the response to the word condition is not shown for frontal language and MD regions. Each fROI shows selective responses to the conditions of interest vs. other conditions across fMRI experiments (temporal lang: Engl > others: p=0.0012, W=98; Frontal lang: p=0.052, W=79; Speech: Nonsn, Engl > others: p<0.0001, W=105; A1: Engl, Nonsn,Txtre > others: p<0.0001, W=105; MD: Hard > others: p=0.067, W=77; Faces: faces > others: p<0.0001, W=105; Objects: Objects > others: p<0.0001, W=105; Scenes: scenes > others: p<0.0001, W=105; Words: words > others: p=0.0002, W=78.

**Supplementary Results 3.** We performed a two-way mixed design ANOVA of language regions’ FC with age group (neonate, adult) as the between-group variable and target (VWFA, faces, scenes, objects) as the within-group variable, and the size of the functional parcels as a covariate. We found that even after accounting for size, the main effects of target (F(3,311) = 7.28, p < 0.001, partial η^2^ = 0.07), group (F(1,311) = 7.49, p = 0.007, partial η^2^ = 0.02), and the interaction (F(3, 311) = 5.65, p = 0.001, partial η^2^ = 0.05) were still significant. We also performed a two-way mixed design ANOVA with age group (neonate, adult) as the between-group variable and target (language regions, multiple-demand regions, speech, A1) as the within-group variable, and the size of the functional parcels as a covariate. Our results showed no main effect of age group (F(1,311) = 3.37, p = 0.067, partial η^2^ = 0.01), significant main effect of target (F(3,311) = 3.1, p = 0.027, partial η^2^ = 0.03), and no significant interaction (F(3,311) = 0.68, p = 0.566, partial η^2^ = 0.01).

**Supplementary Results 4.** We broke up the language regions into frontal and temporal components and performed the same FC analyses as in the main paper but comparing VWFA connectivity to the target regions grouped by temporal and frontal. We found that the VWFA was more connected with temporal language regions than speech and A1 regions in both neonates (Speech: t(39) = 6.22, p < 0.001, Cohen’s d = 0.98, corrected, 95% CI = [0.08, 0.16]; A1: t(39) = 6.01, p < 0.001, Cohen’s d = 0.95, corrected, 95% CI = [0.11, 0.22]) and adults (Speech: t(39) = 5.48, p < 0.001, Cohen’s d = 0.87, corrected, 95% CI = [0.09, 0.20]; A1: t(39) = 7.16, p < 0.001, Cohen’s d = 1.13, corrected, 95% CI = [0.17, 0.31]; Supplementary Figure 3a). We also found that the VWFA connected more with frontal language regions than MD regions in both neonates (t(39) = 4.51, p < 0.001; Cohen’s d = 0.71, corrected, 95% CI = [0.05, 0.13]) and adults (t(39) = 9.71, p < 0.001, Cohen’s d = 1.54, corrected, 95% CI = [0.23, 0.35]; Supplementary Figure 3b).

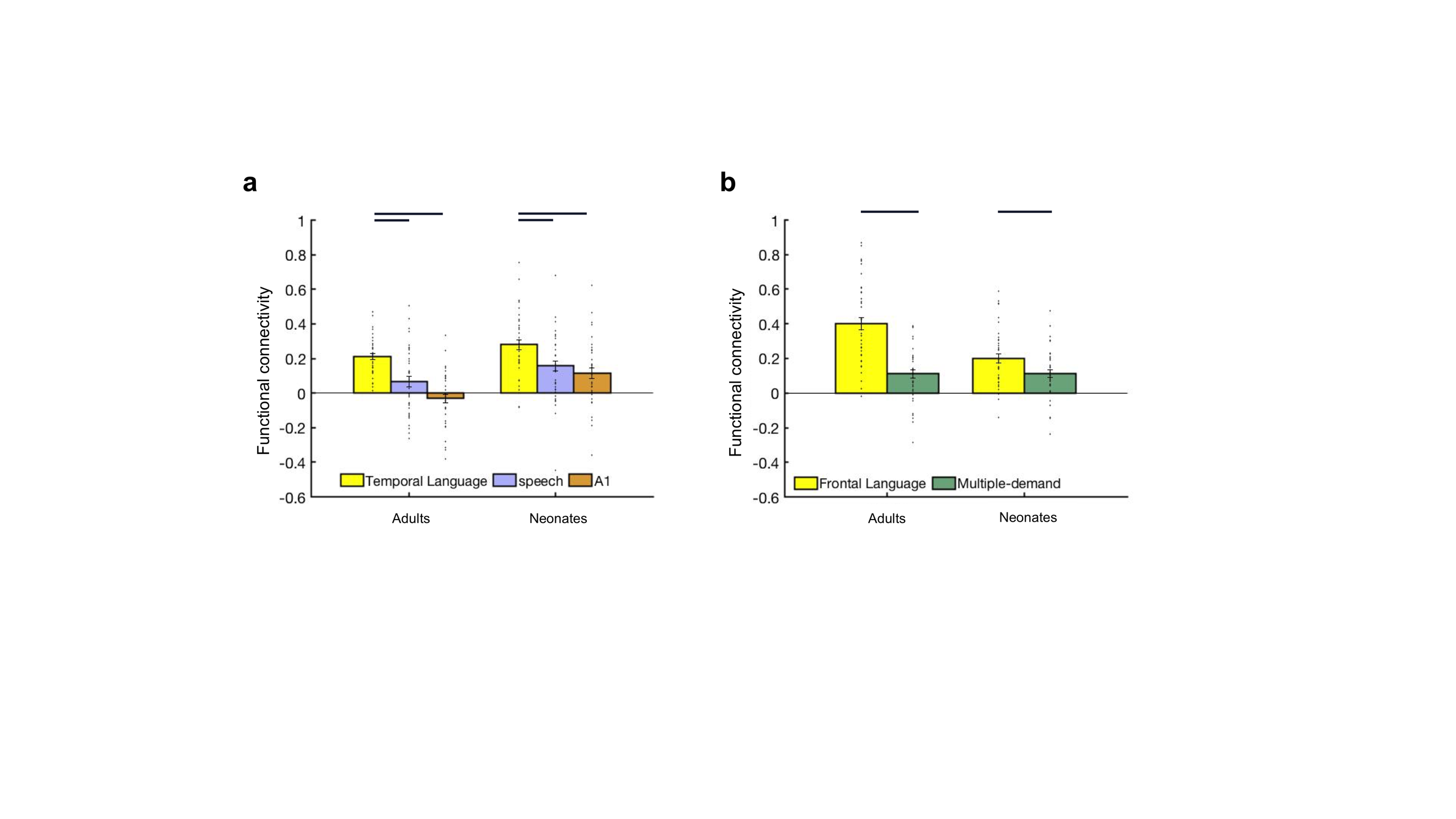

**Supplementary Figure 3 | Functional connectivity between VWFA (seed) and temporal and frontal regions. (a)** Mean FC between VWFA and regions in temporal (i.e., temporal language regions, speech, and A1). **(b)** Mean FC between VWFA and regions in frontal cortex (i.e., frontal language regions, multiple-demand regions). Connectivity values were Fisher z transformed. Error bars denote s.e.m. Individual data points (n = 40 for each age group) were shown for each category. Horizontal bars reflect significant post hoc paired t-tests p < 0.05, corrected.

We also performed the same parametric voxelwise analyses as done for the VTC in the main paper, but here we used the VWFA as the seed and characterized its FC to temporal and frontal cortex. Consistent with parcel-based analyses, when parametrically increasing threshold of FCs, we found that voxels revealed higher FC to VWFA located in temporal and frontal cortex that happen to be in language regions (vs. Speech and A1 regions in temporal, Supplementary Figure 4a; and vs. MD regions in frontal, Supplementary Figure 4b; heatmaps for VWFA’s FC in frontotemporal cortices were also shown on representative adults and neonates surfaces, Supplementary Figure 4c).

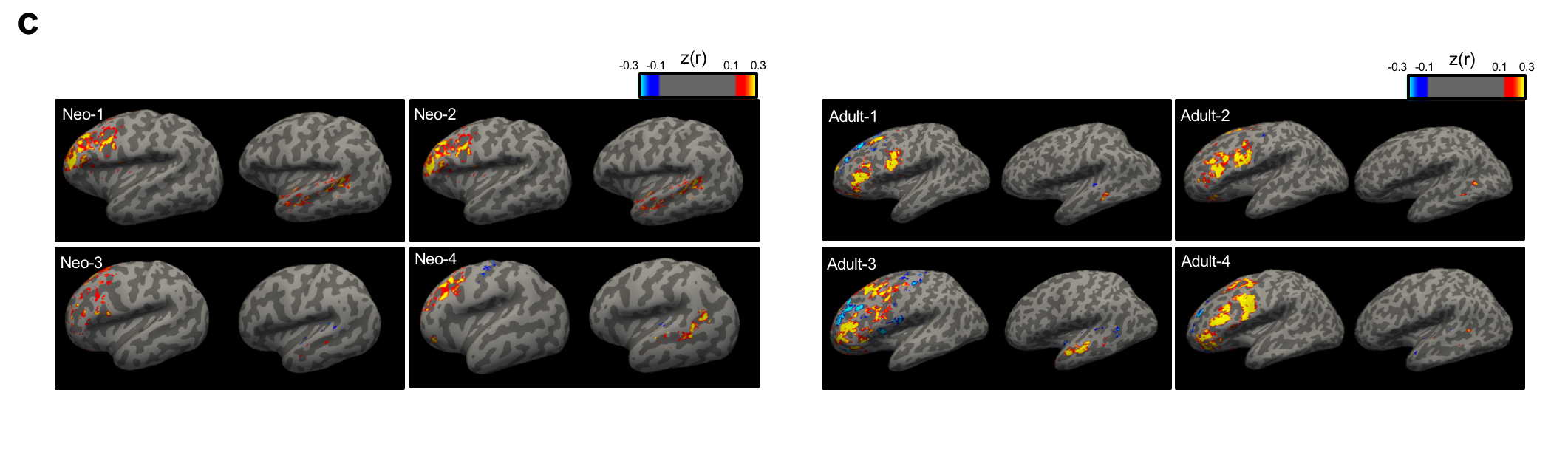

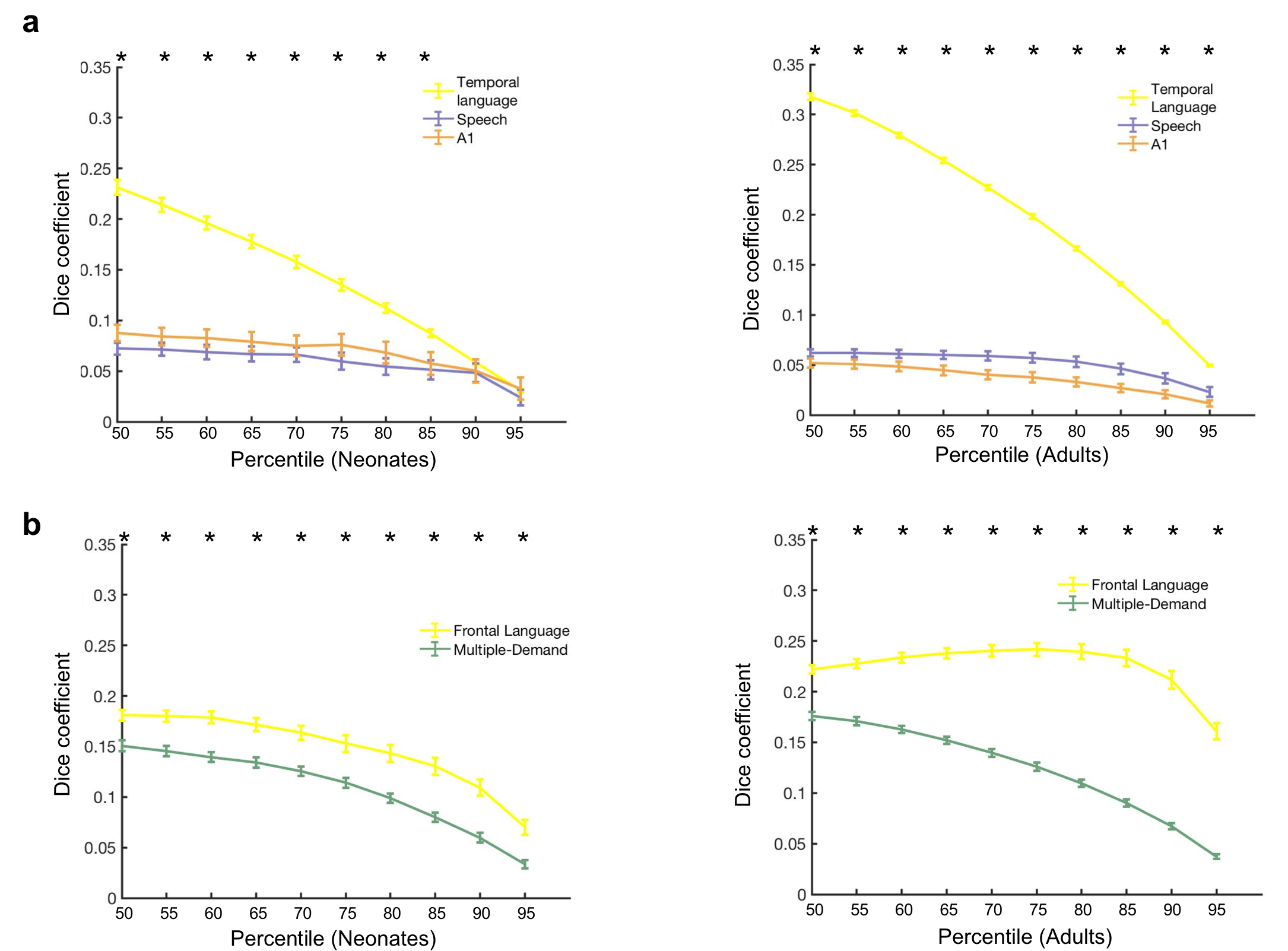

**Supplementary Figure 4 | Voxel-wise analyses from VWFA to frontotemporal cortices. (a)** As we parametrically increasing the threshold of FC within temporal cortex, we quantified how many of these voxels belonged in each functional category using *Dice coefficient*. Averaged FC (Fisher’s z transformed) across neonates (n = 40; 50^th^: z(r) = 0.19, p < 0.001; 95^th^: z(r) = 0.39, p < 0.001); Average FC across adults (n = 40; 50^th^: z(r) = 0.17, p < 0.001; 95^th^: z(r) = 0.42, p < 0.001). **(b)** As we parametrically increasing the threshold of FC within frontal cortex, we quantified how many of these voxels belonged in each functional category using Dice coefficient. Averaged FC across neonates (n = 40; 50^th^: z(r) = 0.22, p < 0.001; 95^th^: z(r) = 0.40, p < 0.001); Average FC across adults (n = 40; 50^th^: z(r) = 0.16, p < 0.001; 95^th^: z(r) = 0.39, p < 0.001). * denotes significant paired t-test (temporal/frontal language vs. averaged of other adjacent functional regions, p < 0.05, corrected). **(c)** Heatmaps for VWFA’s connectivity within frontotemporal in representative neonates and adults, thresholded at z(r) greater than 0.1 (p < 0.001).

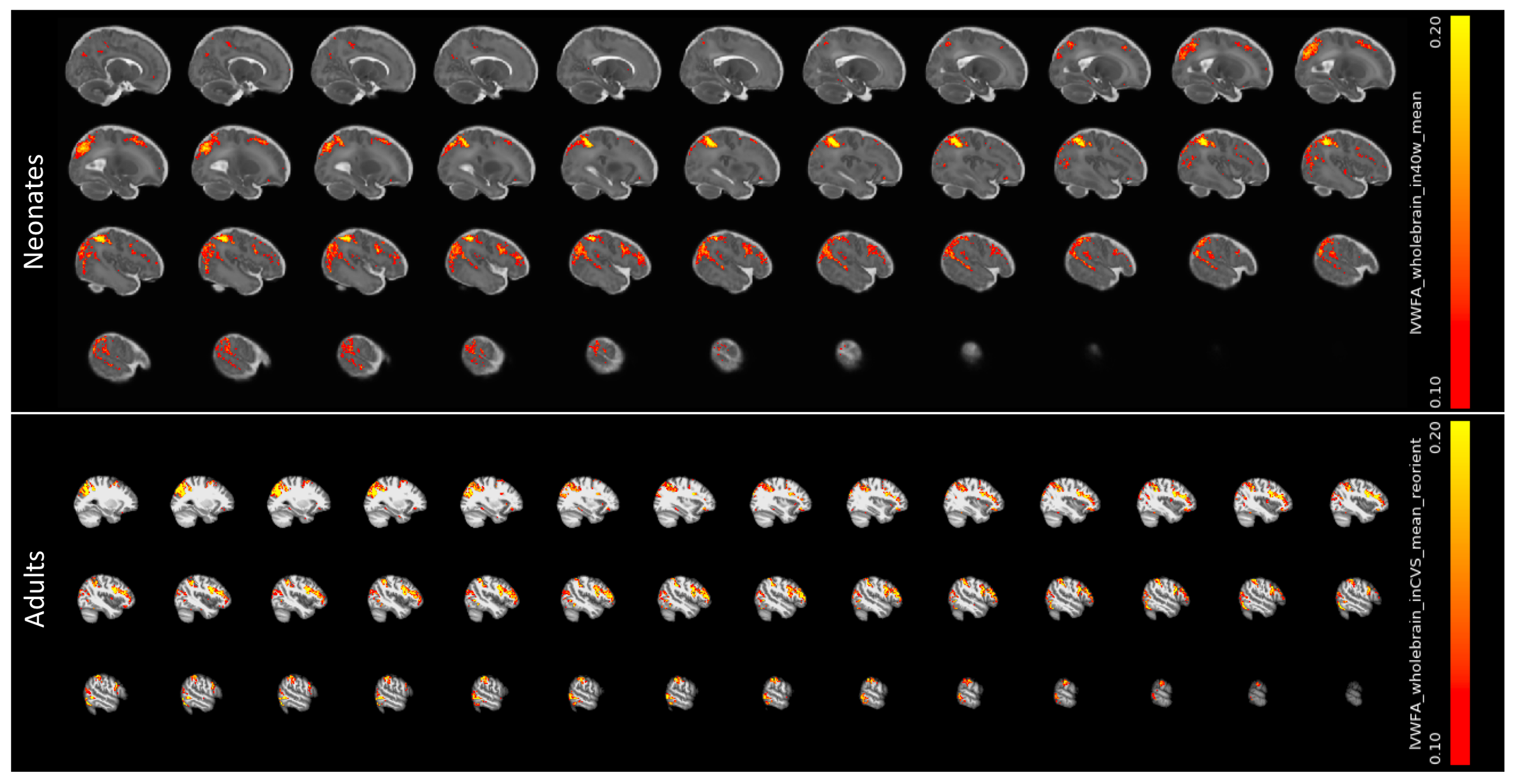

**Supplementary Figure 5 | Averaged whole-brain functional connectivity maps of VWFA.** VWFA whole-brain functional connectivity map was calculated (fisher’s z transformed, z(r)) for each individual. All adults’ maps were register to Freesurfer CVS_avg35_inMNI152 template brain, and all neonates’ maps were registered to 40-week template brain (Serag et al., 2012). The resultant averaged maps were threshold at 0.1< z(r) <0.3.

**Supplementary Results 5.** We obtained association test maps from a meta-analysis approach (Neurosynth, [https://neurosynth.org](https://neurosynth.org/)) for all functions of interest (terms used: language (*1101 studies*), speech (*642*), primary auditory (*114*), faces (*864*), objects (*692*), place (*189*), visual word (*117*)) with a threshold of z > 3, and took the intersection between the functional parcels used in the main analyses with these Neurosynth-generated maps. These maps shared similar spatial locations with our functional parcels, but yield more conservative regions for most of functional categories (Supplementary Figure 6a and 6b, also see Supplementary Table 1 for sizes of these Neurosynth-overlapped parcels). Our main results remain the same with these new parcels (Supplementary Figure 6c-6e): language regions showed significantly higher connectivity to VWFA compared to other adjacent visual regions in adults (Faces: t(39) = 8.46, p < 0.001; Scenes: t(39) = 8.82, p < 0.001; Objects: t(39) = 10.34, p < 0.001, corrected). Similarly, neonates’ language regions also connected more to VWFA than to faces (t(39) = 7.11, p < 0.001, corrected) and scenes (t(39) = 7.19, p = 0.001, corrected) but not objects (t(39) = 1.98, p = 0.054). When examining whether VWFA connects more to language regions to vicinity of language areas, we found that VWFA showed higher FC to temporal language regions than to speech (adults: t(39) = 5.36, p < 0.001, Cohen’s d = 0.85, 95% CI = [0.08, 0.17]; neonates: t(39) = 3.64, p = 0.001, Cohen’s d = 0.58, 95% CI = [0.03, 0.11],corrected) and A1 (adults: t(39) = 6.95, p < 0.001, Cohen’s d = 1.10, 95% CI = [0.16, 0.29]; neonates: t(39) = 3.94, p < 0.001, Cohen’s d = 0.62, 95% CI = [0.05, 0.16], corrected) and higher FC to frontal language regions to adjacent MD regions (adults: t(39) = 4.47, p < 0.001, Cohen’s d = 0.71, 95% CI = [0.05, 0.16]; neonates: t(39) = 4.46, p < 0.001, Cohen’s d = 0.71, 95% CI = [0.05, 0.14]) in both adults and neonates.

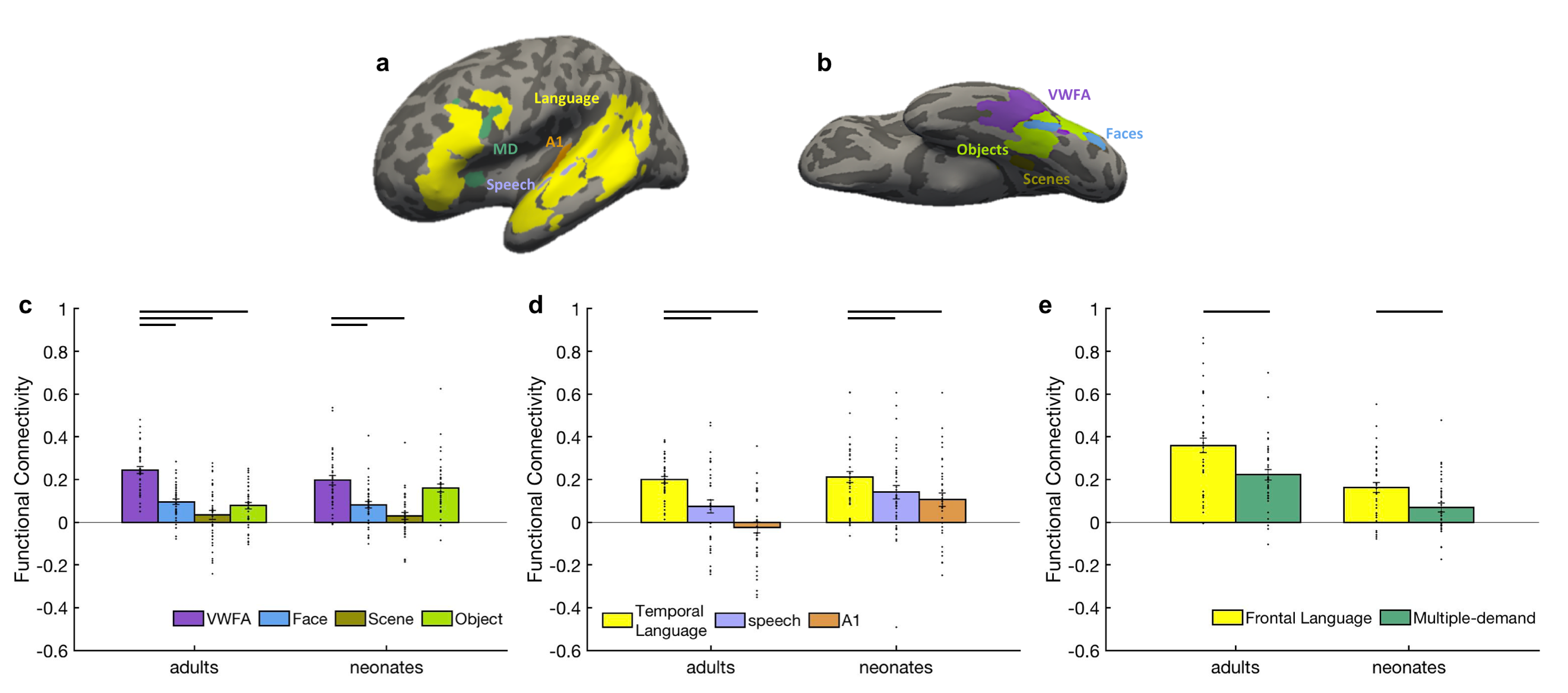

**Supplementary Figure 6 | Functional connectivity results using new regions that intersected with Neurosynth-generated functional maps. (a)** New parcels in temporal and frontal cortex. **(b)** New parcels in ventral temporal cortex. **(c)** Mean FC between language regions and ventral temporal visual regions (i.e., VWFA, faces, objects and scenes). **(d)** Mean FC between VWFA and regions in temporal cortex (i.e., temporal language regions, speech and A1)). **(e)** Mean FC between VWFA and regions in frontal cortex (i.e., frontal language regions, multiple-demand regions). Connectivity values were Fisher z transformed. Error bars denote s.e.m. Individual data points (n = 40 for each age group) were shown for each category. Horizontal bars reflect significant post hoc paired t-tests p < 0.05, corrected.

**Supplementary Results 6.** We used ANTs to register parcels to neonates and adults because it’s commonly used in developmental studies (Avants et al., 2014; Menze et al., 2014; Wang & Yushkevich, 2013). We manually checked the registration results to ensure that functional parcels were in the relatively right locations (see Supplementary Figure 7a for example registration output for a representative adult and neonate). Next, for a quantitative analysis, we compared our registration accuracy to published registration methods used in a recent and comparable neuroimaging study of infants, where functional parcels were again defined based on fMRI data in a group of adults (and in fact using the same atlas/parcels that we are using here), and then registered to the infant brain but using FLIRT (FMRIB's Linear Image Registration Tool from FSL) instead of ANTs (Deen et al., 2017; Kamps et al., 2020). We took the binary gray matter mask from CVS average-35 MNI152 template brain and then registered it to individual’s native space with both ANTs and FLIRT. The registration result was compared to each individual’s own binary gray matter mask. We found that in general, both methods yield high accuracy (over 90%), but ANTs significantly outperformed FLIRT (t(39)=6.21, p < 0.001, Cohen’s d=0.62, 95% CI=[0.006, 0.01]; Supplementary Figure 7b).

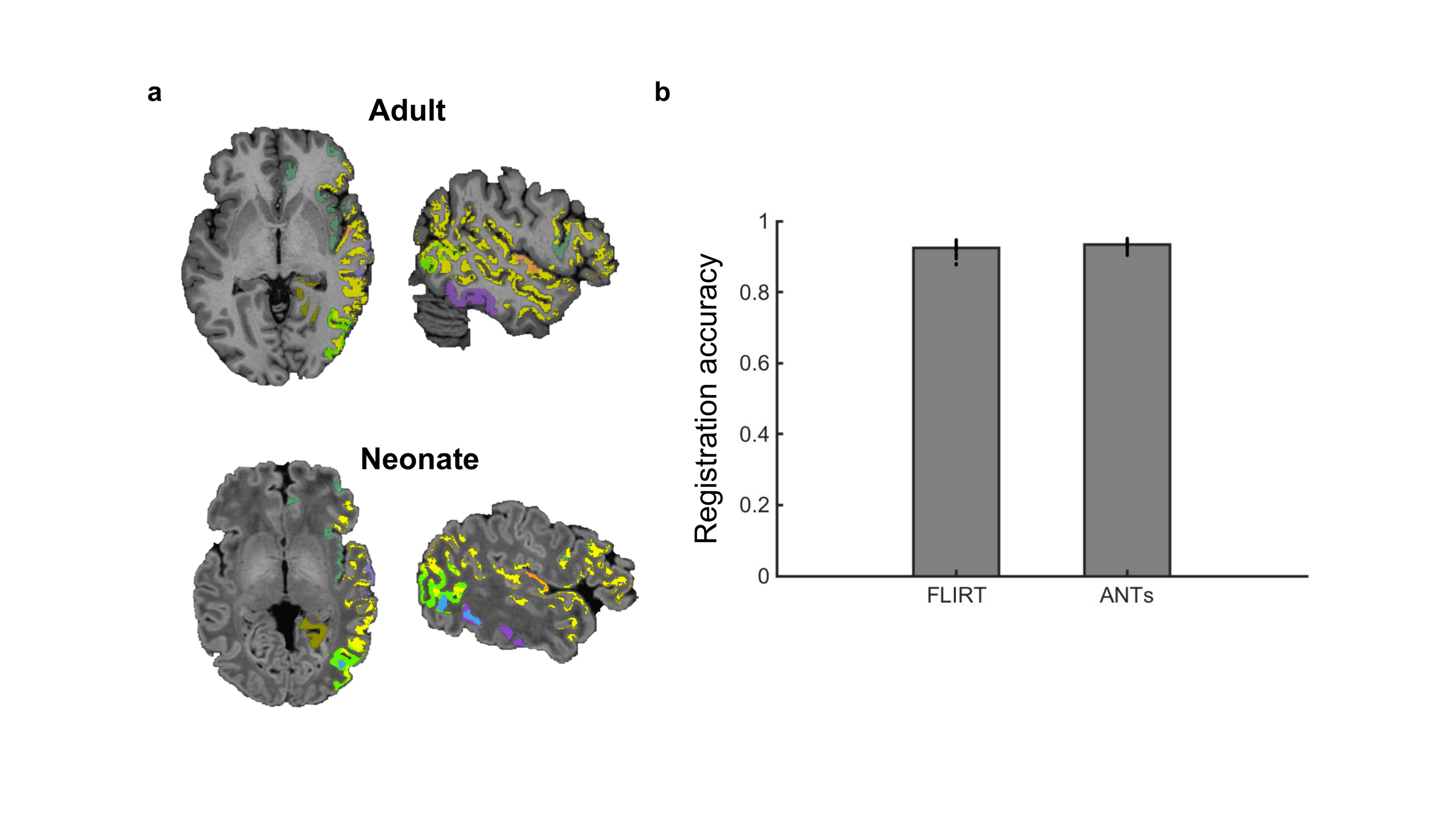

**Language speech A1**

**MD**

**VWFA Faces Scenes Objects**

**Supplementary Figure 7 | Registration results.** **(a)** Registration results for all parcels from 8 categories for a representative adult (top) and neonate (bottom). **(b)** Comparing registration accuracy of binary gray matter mask using FLIRT and ANTs. ANTs significantly outperformed FLIRT (n = 40; t(39)=6.21, p < 0.001, Cohen’s d = 0.62, 95% CI = [0.006, 0.01]).

**Supplementary Table 1. Information of parcels used in the current study.**

| Category | How to define | Regions (Left) | Coordinate  (CVS_avg35_inMNI152 template space) | | | Size in mm^3^ (original/within Neurosynth) | | Source (number of subjects) |
| --- | --- | --- | --- | --- | --- | --- | --- | --- |
|  |  |  |  |  |  | Adults | Neonates |  |
| Words | words vs. objects | VWFA | 174 | 165 | 99 | 4688/3632 | 1163/815 | Saygin et al., 2016 (20) |
| Face | Faces vs. objects | FFA | 168 | 162 | 90 | 352/312 | 99/80 | Julian et al., 2012 (40) |
|  |  | OFA | 164 | 154 | 65 |  |  |  |
| Scene | Scenes vs. obj | PPA | 149 | 151 | 94 | 2552/536 | 616/60 |  |
| Object | Obj vs. scr.obj | LO PFS | 168 | 140 | 70 | 3608/2560 | 994/626 |  |
|  |  |  | 163 | 161 | 94 |  |  |  |
| Language | Engl vs. nonsn | *Temporal* |  |  |  | 5032/1976 | 845/328 | Fedorenko et al., 2010 (25) |
|  |  | MidPostTemp | 184 | 137 | 107 |  |  |  |
|  |  | PostTemp | 177 | 132 | 83 |  |  |  |
|  |  | MidAntTemp | 184 | 159 | 128 |  |  |  |
|  |  | AntTemp | 179 | 162 | 147 |  |  |  |
|  |  | AngG | 166 | 118 | 69 |  |  |  |
|  |  | *Frontal* |  |  |  | 4016/2648 | 676/427 |  |
|  |  | IFG | 174 | 123 | 161 |  |  |  |
|  |  | IFGorb | 169 | 149 | 182 |  |  |  |
| Multiple-demand | Hard vs. easy | SMA | 134 | 87 | 145 | 2760/336 | 388/40 | Fedorenko et al., 2013 (40) |
|  |  | ACC | 134 | 134 | 177 |  |  |  |
|  |  | IFGorb | 175 | 132 | 154 |  |  |  |
|  |  | MFG | 161 | 115 | 178 |  |  |  |
|  |  | MFGorb | 159 | 156 | 195 |  |  |  |
|  |  | Insula | 161 | 145 | 148 |  |  |  |
| Speech | bisyllabic pseudowords vs. baseline | within STG | 189 | 149 | 126 | 1256/1256 | 209/209 | Basilakos et al., 2018 (20) |
| A1 | Anatomically defined | Primary auditory cortex | 173 | 140 | 127 | 744/704 | 119/109 | Desikan et al., 2006 |

* Engl, English sentences; nonsn, non-word sentences; Obj, objects; src.obj, scrambled objects; VWFA, visual word form area; FFA, fusiform face area; OFA, occipital face area; PPA, parahippocampal place area; LO, lateral occipital; PFS, posterior fusiform sulcus; AntTemp, anterior temporal lobe; MidAntTemp, middle-anterior temporal lobe; MidPostTemp, middle-posterior temporal lobe; PostTemp, posterior temporal lobe; AngG, angular gyrus; IFG, interior frontal gyrus; IFGorb, orbital IFG; MFGorb, orbital part of the middle frontal gyrus, IFGop, opercular part of the inferior frontal gyrus; SMA, supplementary motor area; ACC, anterior/mid cingulate cortex; STG, superior temporal gyrus.
